# Exploring transient neurophysiological states through local and time-varying measures of Information Dynamics

**DOI:** 10.1101/2024.06.19.599743

**Authors:** Y. Antonacci, C. Barà, G. De Felice, A. Sferlazza, R. Pernice, L. Faes

## Abstract

Studying the temporal evolution of complex systems requires tools able to detect the presence and quantify the strength of predictable dynamics within their output signals. Information theory aids in such a description, particularly through information storage (IS), which reflects the regularity of system dynamics by measuring the information shared between the present and the past system states. While the conventional IS computation provides an overall measure of predictable information, transient behaviors of predictability occurring during system transitions can be assessed by time-resolved measures such as the local information storage (L-IS), assuming stationarity, and the time-varying information storage (TV-IS), without stationarity assumptions. In this work, through a comparative analysis in simulated and real contexts, we aim to demonstrate how these methods complement each other and reveal dynamic changes of the system behavior associated to state transitions. In simulations, we show that the TV-IS can effectively track sudden changes of the information stored in the system, which is reflected in its average value computed over specific time intervals; on the other hand, the surprise originated by the emergence of a change in the predictability of the system is reflected in the variance of the L-IS computed within specific time intervals. In neurophysiological applications, the distinct phenomena of respiratory activity in sleep apnea and brain activity during somatosensory stimulation both reveal a significant decrease of IS evoked by state transitions, highlighting how such transitions can inject new information in physiological systems, affecting significantly their internal dynamics. Overall, TV-IS and L-IS appear to provide different and complementary information about the behavior of the systems under investigation, thereby offering valuable tools for the study of complex physiological systems where both stationary and non-stationary conditions may be present.

## I. INTRODUCTION

In recent decades, the study of complex systems has been significantly advanced by the exploration of deterministic and probabilistic approaches to time series analysis. These methodologies offer distinct perspectives on understanding system behavior within diverse physical, biological, and technological contexts. Deterministic approaches, grounded in precise mathematical formulations, aim to unveil the underlying dynamics governing the evolution over time of the observed system. In contrast, probabilistic approaches aim to characterize such dynamics via descriptors derived from the statistical analysis of the stochastic processes assumed to map the system evolution. Interestingly, probabilistic approaches can be undertaken also in a deterministic framework, when the observed dynamics are complex enough (e.g., chaotic) to retain some degree of unpredictability [1]. In this context, an unified description of complex systems comes from the field of information theory, which offers general and flexible tools for modeling the system dynamics and inferring their emergent properties [2, 3].

Turing suggested that every act of information processing within a network of interacting systems can be decomposed into three distinct processes: information storage, information transfer, and information modification [4]. In particular, information storage (IS) is the main information-theoretic concept that is exploited to describe the dynamics of an individual units of a complex system from an observed time series. IS can be defined as the information content of the past states of a dynamic system that can be used to predict the current state [5]. Thus, it allows to measure the regularity and predictability of a dynamic process and, under the Gaussian assumption, can be estimated with the identification of a simple linear model [6, 7]. IS has found wide applications in various fields, including the study of cerebrovascular and cardiorespiratory dynamics [8] and the distribution of neural information processing [9].

The standard computation of IS lacks the capability to provide time-resolved information about the dynamics of the analyzed system. Nevertheless, capturing transient behaviors of dynamic processes can provide new insights into how a system adapts to changes and perturbations, which is crucial in several contexts. To this end, time-resolved measures of IS have been recently proposed to identify and analyze transiently changing properties of dynamical systems, marking for instance the transition across different states. The so-called local information storage (L-IS) investigates the predictability and regularity of the system’s dynamics at each time instant [5, 9, 10]. The L-IS can be computed from the parameters of a linear auto-regressive (AR) model identified on an observed time series under the assumption of stationarity. An alternative is represented by the time-varying IS (TV-IS) [11] which, unlike the local approach [10], does not require stationary signals. It is based on a recursive version of the least squares analysis called recursive least squares (RLS), which involves a time-varying identification of a linear AR model [12] and has proven to be a reliable way to investigate the time-specific dynamics of a complex system [11, 13, 14].

Although both L-IS and TV-IS have proven to be useful in capturing dynamic changes in the predictable information produced by complex systems, it is not clear what properties of the analyzed transient dynamics they reflect, what is the impact of their different assumptions, to what extent they yield similar information, and how they perform in different scenarios. In this work, through a comparative analysis in simulated and real physiological dynamics, we aim to elucidate how these methodologies complement each other and allow uncovering time-resolved behavior of complex systems. Our focus is on assessing the capability of L-IS and TV-IS to capture transient neurophysiological states, and to highlight their shared and complementary properties. To this end, we first use well-controlled simulated data to reproduce complex dynamics in the framework of linear systems. We then analyze the respiratory dynamics during sleep apneas [15], which are known to induce changes in signal regularity [16], and real benchmark epicranial electroencephalographic (EEG) data recorded during whisker stimulation in rats, where the timing of activation of the different brain regions involved in stimulus encoding is well known [17, 18]. To stimulate the use of these measures in the analysis of neurophysiological signals, the algorithms for the computation of L-IS and TV-IS are made available through the Time Specific Analysis (TSA) Matlab toolbox, freely accessible from https://github.com/YuriAntonacci/TSA toolbox.

## MATERIALS AND METHODS

This section outlines the methodological approaches utilized for assessing the predictability of a random process at each time step *n*, under either stationary or non-stationary conditions. The following subsections describe the techniques employed for estimating local and time-varying predictability measures using a linear autoregressive model.

### A. Information Storage

Let us take into account a dynamic system *Y*, whose progression over time is governed by the stochastic process *Y* = *{Y*_*n*_, *n* ∈ ℤ*}*. Assuming that *Y* is a Markov process with finite memory of order *p*, its whole past history can be truncated using *p* time steps, i.e., using the *p*−dimensional variable *W*_*n*_ = [*Y*_*n*−1_, …, *Y*_*n*−*p*_]^⊤^ ∈ ℝ^*p×*1^. Then, the IS is defined as the information shared between the present and past states of the process and can be computed as follows [5, 9]:

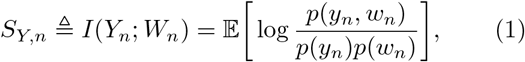

where *I*(·; ·) denotes the mutual information, *p*(·, ·) and *p*(·) denote respectively the joint and the marginal probability density function (PDF), and *y*_*n*_ and *w*_*n*_ refer to realizations of *Y*_*n*_ and *W*_*n*_, respectively. The IS measures the predictability of the process at time *n* by quantifying the average level of uncertainty about the current state of the process (i.e., *Y*_*n*_) that can be resolved by the knowledge of its past states (i.e., *W*_*n*_).

The argument of the expected value reported in (1) represents the so-called local information storage, *s*_*y*,*n*_ = log (*p*(*y*_*n*_, *w*_*n*_)*/p*(*y*_*n*_)*p*(*w*_*n*_)) [5, 9]. Without prior assumptions on the statistical properties of the process *Y*, when multiple realizations are available, it is possible to obtain an estimate of the local information storage *ŝ*_*y*,*n*_ at each time step *n*. On the other hand, if only a single realization of *Y* is available in the form of the finite-length time series *y* = *{y*_1_, …, *y*_*N*_ *}*, two options are available: i) the process *Y* is assumed to be stationary and ergodic: in this case, an estimate of the local IS *ŝ*_*y*,*n*_ can be obtained from the time series *y* and its mean value *Ŝ*_*Y*,*n*_ = *Ŝ*_*Y*_, returns the well-known time invariant IS measure [9], or ii) the process *Y* is assumed to be non-stationary: it is possible to compute *Ŝ*_*Y*,*n*_ with a time-varying approach assuming local stationarity of the dynamics as introduced in [11]. Both possibilities are discussed and analyzed in detail below by exploiting parametric models. Specifically, adopting a joint Gaussian distribution for the variables that constitute the current and past states of the observed process *Y*, both L-IS and TV-IS can be computed starting from the identification of an AR model and its time-variant version respectively [11, 19].

#### 1. Stationarity assumption: local approach

The L-IS represents the amount of stored information used by the process in the specific time instant *n*. Note that, unlike the IS, the L-IS is not bounded and could assume negatives values [5, 9, 10]. In particular, positive values of *s*_*y*,*n*_ occur when knowing the past state *w*_*n*_ increases the probability of observing the present state *y*_*n*_, while negative values of *s*_*y*,*n*_ are measured when the past of the process is misinformative about the current state and thus reduces its probability of occurrence. In the linear parametric framework, the current state *Y*_*n*_ of the process *Y* can be predicted as a linear combination of the past states *W*_*n*_ by means of an AR model: *Y*_*n*_ = *AW*_*n*_ + *U*_*n*_, where *A* = [*a*_1_, …, *a*_*p*_] ∈ ℝ^1*×p*^ is the coefficient vector which, under the assumption of stationarity, is independent of the time *n*, and *U*_*n*_ is the prediction error assumed to be white, with zero mean and with variance 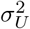. Then, an explicit formulation of the L-IS for Gaussian processes can be written as follows [10]:

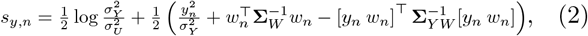

where 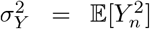 is the variance of 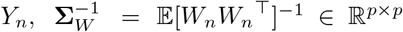 is the inverse of the covariance matrix of *W*_*n*_, and 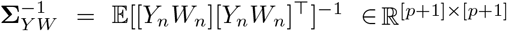 is the inverse of the joint covariance matrix of *Y*_*n*_ and *W*_*n*_. In the Eq. (2), the first term corresponds to the IS (i.e., *S*_*Y*_), while the second term, which changes at any time step *n*, has zero average. Given (2), we can notice that the estimation of the local IS reduces to computing the relevant covariance and cross-covariance matrices between the present and the past variables of the process, i.e., 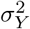,Σ_*W*_, andΣ_*YW*_, together with the innovation variance 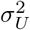. These matrices can be derived from the autocovariance structure of the linear AR representation of the process *Y*. A complete derivation of the above mentioned matrices via the solution of extended Yule-Walker equations together with a detailed description of the use of linear models exploited to estimate L-IS can be found in [7, 10]. Note that, in practical applications, the covariance matrices in (2) are computed under the hypothesis of ergodicity of the process *Y* from a single realization available in the form of the finite-length time series *y* = *{y*_1_, …, *y*_*N*_ *}*.

#### 2. Non-stationarity assumption: time-varying approach

Relaxing the stationarity assumption for the process *Y*, its current state *Y*_*n*_ can be predicted as a linear combination of its past state *W*_*n*_ by means of a time-varying version (TV-AR) of the AR model defined in the previous section: 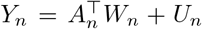, where *U*_*n*_ is the prediction error and *A*_*n*_ = [*a*_1,*n*_, …, *a*_*p*,*n*_]^⊤^ ∈ ℝ^*p×*1^ is the vector of the TV AR coefficients that need to be estimated at each time step *n*. Under the joint Gaussian assumption for *Y*_*n*_ and *W*_*n*_, the relation stated in Eq. (1) can be written as [11]:

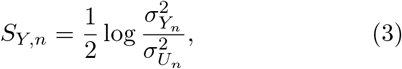

where 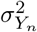 and 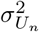 are respectively the variance of the process *Y* and the variance of the prediction error *U* sampled at the time instant *n*.

The estimation procedure of the TV-IS can be performed through a recursive implementation of the least squares method [11, 20]. The RLS method consists in the following computation steps: (i) choose a value for the adaption factor *c* ∈ (0, 1) and an order *p* of the AR model; (ii) define proper initial conditions for the vector of coefficients at time *p, A*_*p*_ = [*a*_1,*p*_, …, *a*_*p*,*p*_]^⊤^ ∈ ℝ ^*p×*1^ and for the correlation matrix of the past state of *Y* stored in *W*_*p*_, 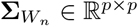; (iii) considering *N-p* consecutive time steps, for *n* = *p* + 1 to *N* repeat the following steps:

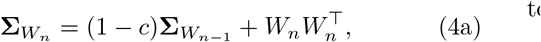

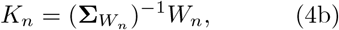

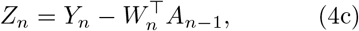

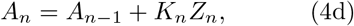

where *K*_*n*_ ∈ ℝ ^*p×*1^ is the so-called gain vector and *Z*_*n*_ ∈ ℝ ^1*×*1^ is intended as the *a-priori* estimation error that can be viewed as a “tentative” value of the error before updating the AR coefficients vector. The parameter *c* ensures that the data in the distant past are “forgetten” in order to afford the possibility of following the statistical variations in non-stationary conditions. For a detailed mathematical derivation of the RLS solution we refer to the supplementary material of [11]. Following the results of [21], a recursive estimation of the time-varying innovation variance can be obtained as follows: 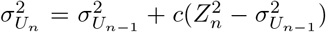. Moreover, to complete the computation of the TV-IS as in Eq. (3), a recursive estimation of the process variance 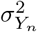can be directly derived from the structure of the linear TV-AR representation of the process *Y* [11].

We note that under the assumption of stationarity of the process *Y*, a connection between the local and the time-specific IS definitions can be drawn. In particular, the expected value of the L-IS as defined in Eq. (2) corresponds to the time-specific IS defined in Eq. (3), i.e., E[*s*_*y*,*n*_] = *S*_*Y*,*n*_ = *S*_*Y*_. This is ensured by the invariance over time of the statistical moments of a stationary AR process (i.e., 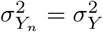 and 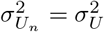).

## II. SIMULATION STUDY

In this section we explore the behaviour of both L-IS and TV-IS on simulated linear AR dynamics obtained by modifying the statistical structure of the process over time. Specifically, we design a first-order TV univariate AR process defined as *Y*_*n*_ = *a*_1,*n*_*Y*_*n*−1_ + *U*_*n*_, where *U*_*n*_ is a vector of zero-mean white Gaussian noise with variance 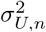, and *a*_1,*n*_ is the coupling strength between the past and the present states of the process *Y*, controlling the information stored in the system at each time instant *n*. In particular, high values of *a*_1,*n*_ indicate a greater amount of information stored in the system and thus, higher predictability of the system dynamics. One realization of *N* = 60 · 10^3^ samples was generated while *a*_1,*n*_ and *σ*^2^ were set to vary over time with the pre-determined structures, reported in Figure 1, and combined to simulate three different scenarios. Specifically, both the autoregressive coefficient *a*_1,*n*_ and the noise variance 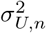 were enabled to vary according to two different waveforms, i.e., a constant value equal to 0.8 and a periodic square waveform alternating between 0.3 and 0.9 with a duty cycle of 50%. For all the analyzed cases, the forgetting factor for RLS was set to 1 − *c* = 0.98 following the results of previous works [13**?** ] while the model order *p* was set to 1.

**FIG. 1.**
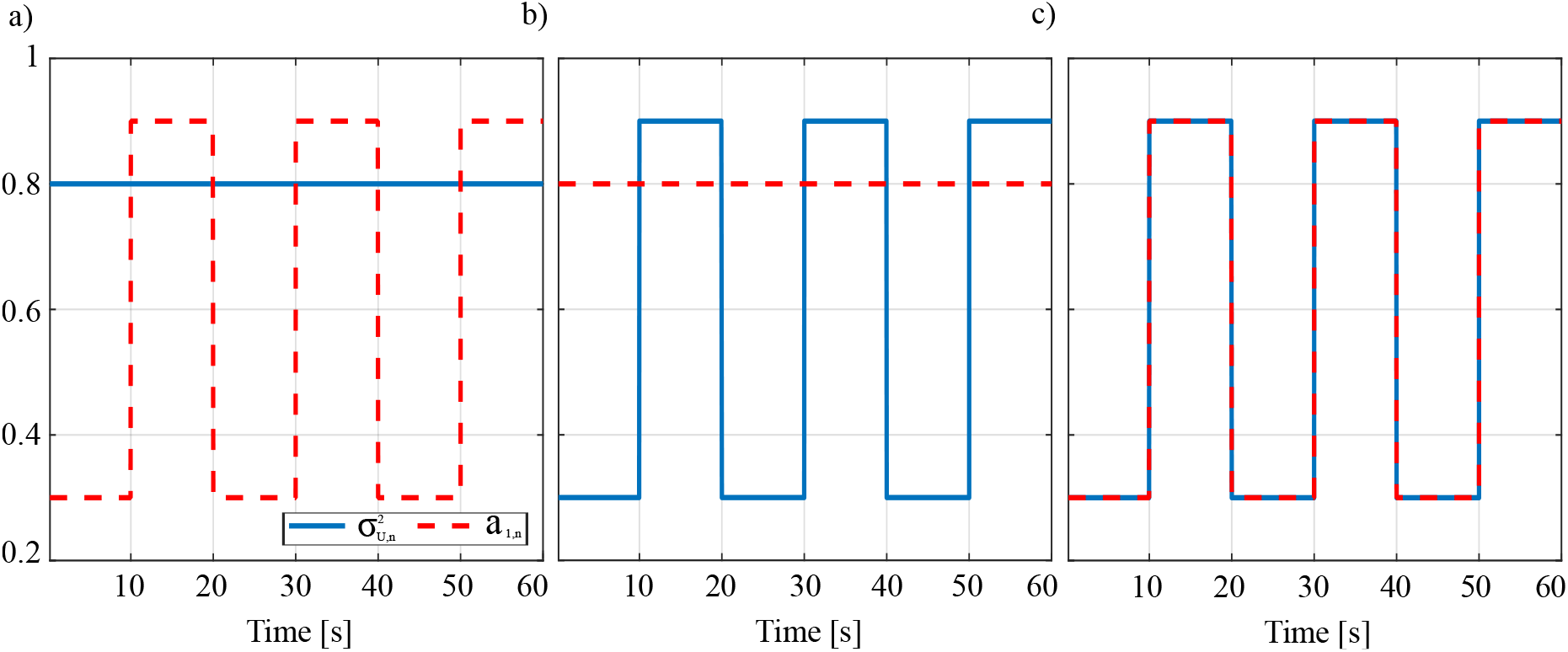
Evolution over time of the autoregressive coefficient *a*_1,*n*_ (dashed red line) and the noise variance 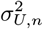 (blue line) for the three different simulated conditions: (i) 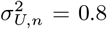 and *a*_1,*n*_ varying according with a square waveform (*a*); (ii) coupling strength remains constant while the noise variance oscillates (*b*); (iii) the noise variance and the coupling strength vary with the same periodic square wave (*c*)

The local and TV approaches for the time-resolved estimation of the IS were applied, respectively assuming the overall stationarity of the simulated process (by considering the several repetition of the alternating pattern in Figure 1) or its local non-stationarity (by considering single instances of the transitions). The choice of one of the two perspectives determines which aspects of system behavior are emphasized and provides a comprehensive understanding of its dynamics.

Figure 2 displays the theoretical and estimated trends of the local and time-varying IS for three different simulated scenarios. The theoretical values of information storage (black line) show that the temporal evolution of IS is not dependent on the modification of the innovation term 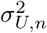, but rather on the value of *a*_1,*n*_ (panels *a* and *d*, and panels *c* and *f*). When only the variance of the innovation term oscillates, as in panel *b* and *e*, the theoretical IS remains constant. This can be explained by the fact that the IS is defined as the ratio between the process variance and the innovation variance (see Eq.(3)). Indeed, since in the second scenario, the coefficient *a*_1,*n*_ does not vary over time, changes in the numerator term follow those in the denominator term, resulting in a value of IS equal to 0.51 at each time step.

**FIG. 2.**
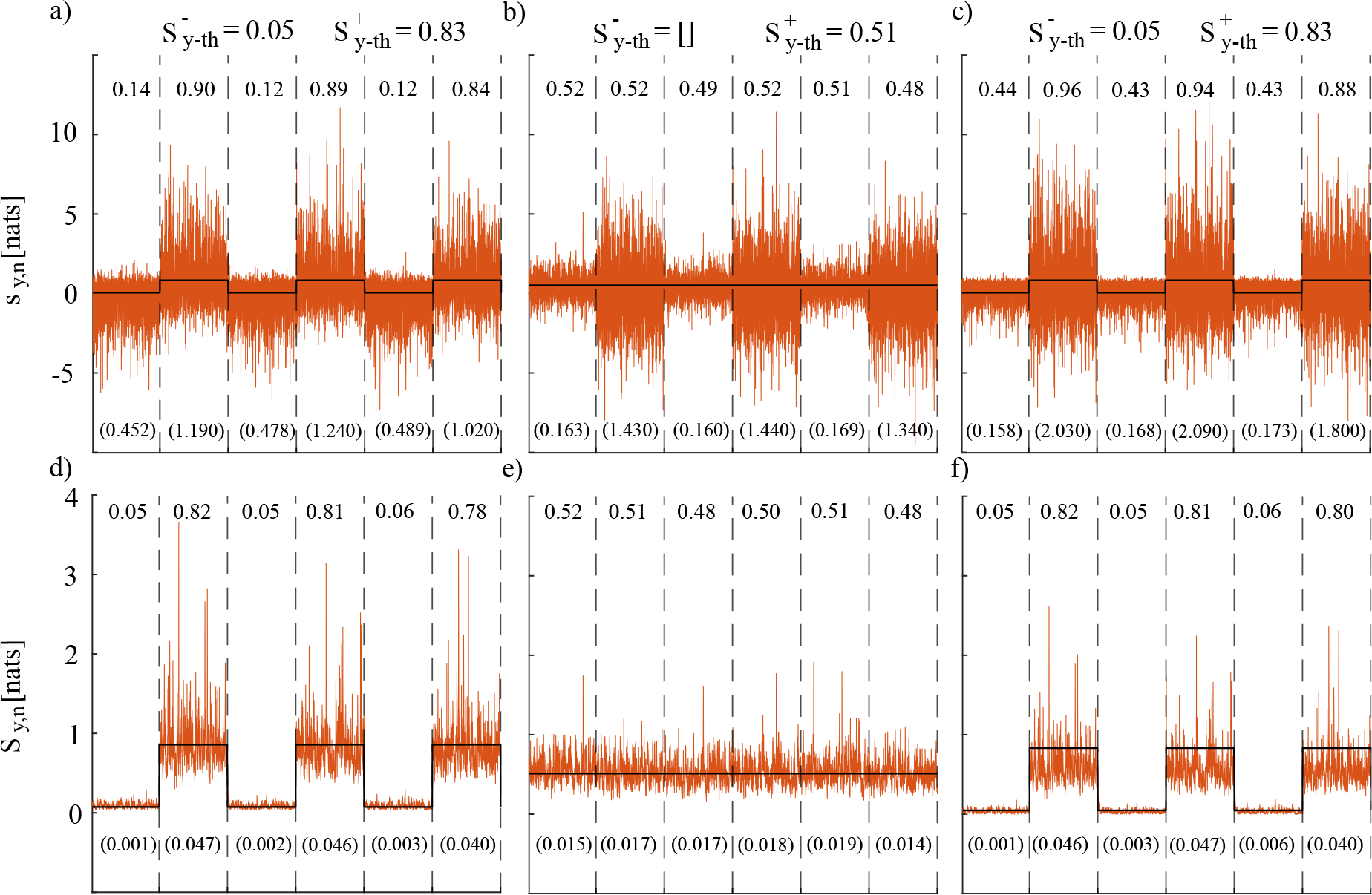
Estimated and theoretical trends (orange and black lines, respectively) for the L-IS (first row) and the TV-IS computed with RLS approach (second row) for the three different simulation settings: (i) constant trend for 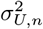 with *a*_1,*n*_ allowed to vary over time (panels *a* and *d*); (ii) constant trend for *a*_1,*n*_ with 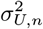 allowed to vary over time (panels *b* and *e*); (iii) variation over time of both 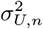 and *a*_1,*n*_ (panels *c* and *f*). At the top of the figure are reported the theoretical values of IS for the highest and lowest steady states (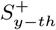 and 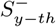). Mean and variance of the estimated values of IS in each time window, delimited by vertical dotted lines, are reported respectively on the top and on the bottom (in brackets) of the figures.

The profiles of the local IS reported in Figures 2 (panels *a*-*c*) document how this measure is sensitive to both the internal dynamics and the unpredictable dynamics of the simulated process. Specifically, when the variability over time is only in the innovation variance (panel *b*), L-IS averaged within each time time interval remains quite close to the theoretical one whereas the profile of its variance in the same time intervals displays a modulation which follows the theoretical innovation variance profile. Otherwise, if the modifications are associated to the coupling strength (panels *a* and *c*), both the mean and the variance of the L-IS computed over each time interval exhibit a modification which correspond to *a*_1,*n*_ modulation. Thus, while the mean L-IS is related to the predictable internal dynamics quantified by the theoretical profile of the time-variant information storage, its variability is modulated both by the internal dynamics and by the unpredictable dynamics quantified by the innovation variance. As regards the nature of the L-IS variability, our results reveal that the L-IS exhibits oscillations between positive and negative values, with the latter being observed when the noise pulls the system in the opposite direction of the coupling between *Y*_*n*_ and *Y*_*n*−1_, making the knowledge of *Y*_*n*−1_ misleading about *Y*_*n*_. On the other hand, large positive values are displayed when the noise and the strength of *a*_1,*n*_ act in the same direction. It is noteworthy that the fluctuations of the L-IS profiles do not merely replicate the modification over time of the innovation variance, but instead represent the interplay between this latter and the trend of *a*_1,*n*_. Indeed, in addition to the mean value, the L-IS variability changes according to the local modifications of the process dynamic. These results are consistent with a previous study that obtained a formulation for local Granger causality in the context of VAR modeling of stationary Gaussian processes [16].

As regard the time-varying approach, the trends depicted in Figure 2 (panels *d-f*) demonstrate that the values of the estimated TV-IS are consistently close to the theoretical ones for each simulated scenario. Moreover, the variance of the TV-IS estimates changes with the strength of the internal dynamics of the system and thus with the TV-IS value itself, but not with the strength of the unpredictable dynamics modulated by 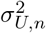. Our findings emphasize the ability of the RLS algorithm to accurately track the transitions between the two imposed states in the system, in agreement to previous studies in the literature that highlighted the accuracy, consistency, and efficiency of RLS for the estimation of time-varying versions of information storage [11], coherence [13], directed transfer function, partial directed coherence [14], and Granger causality [20, 22]. As regards the forgetting factor, previous works have highlighted how the choice of this parameter can affect the estimation performance identifying the range of values between 0.96 and 0.99 to ensure a proper trade-off between speed adaptation to transitions and estimation variance [11, 13, 14, 20, 22]. Our decision to settle on a forgetting factor of 0.98 aligns with the range established in the literature, achieving a balanced trade-off between bias and variance.

Overall, the results of the simulation studies show that TV-IS reliably follows the theoretical value of IS and any deviation from the theoretical value can considered as a prediction error. Thus, the variance observed in TV-IS is primarily imputable to inaccuracies of the RLS estimator rather than to the information content. On the other hand, L-IS often exhibits a less reliable association with the theoretical IS value and, in some instances, it may deviate significant or even fail to follow this latter. Despite this, the variance observed in L-IS serves a different purpose since it reflects the unpredictable dynamics of the analyzed process.

## III. APPLICATIONS TO PHYSIOLOGICAL SYSTEMS

In this section, we explore the effectiveness of local and time-varying IS in detecting state transitions from respiratory data during sleep apnea and from evoked EEG responses during whisker stimulation in rats. The analysis of respiratory dynamics and the study of evoked EEG responses have been already explored by using the local [16] and the time-varying approaches [23], respectively. During sleep apnea, the respiratory dynamics are characterized by relatively stable periods alternated with abrupt transitions occurring with a rhythmic repetition of apneic episodes. Similarly, the examination of evoked potentials constitutes a well-controlled domain where several dynamic responses are elicited by external stimuli, exhibiting rapid changes in amplitude at specific time instants after the onset of the stimulation. We therefore investigate the behavior of the two time-resolved IS estimators in the study of both phenomena, considering either the global evolution of multiple repetitions of the system’s oscillatory patterns (assuming the overall stationarity) or single instances of the transitions (assuming local non-stationarity).

### A. Respiratory dynamics during sleep apneas

The respiratory signal amplitude measured from a subject suffering from sleep apneas was used to investigate how local and time-varying approaches work in distinguishing state transitions in breathing dynamics [15, 24]. The investigated data can be found at https://www.physionet.org/content/santa-fe/1.0.0/ and have been used in a previous work to study cardiorespiratory dynamics interaction using local Granger causality [16].

In Figure 3*a*, the 1200-samples time series collected at 2 Hz is depicted and the apnea intervals characterized by the absence of respiratory oscillations are highlighted in grey. In panels *b* and *c*, we report the temporal profiles of the IS measure obtained using the local and time-varying approaches, respectively. For the local approach, the model order *p* was estimated according to the minimum description length criterion (MDL) [25]. As regards the TV approach, the forgetting factor was set sto 0.98 and, as initial conditions (*A*_*p*_,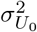) for the RLS algorithm, the result of the identification procedure obtained through an ordinary least squares prediction was used since it was demonstrated to be a good starting point to counteract convergence issues of RLS when few data samples or few realizations are available [26]. One-hundred realizations of time shuffled surrogate data were used to assess point-by-point the statistical significance of the information measures. In particular, at each iteration, a surrogate series was generated through a temporal permutation of samples of the original time series [27]. In this way, any temporal correlation of the signal was destroyed and a null distribution for the IS was obtained.

**FIG. 3.**
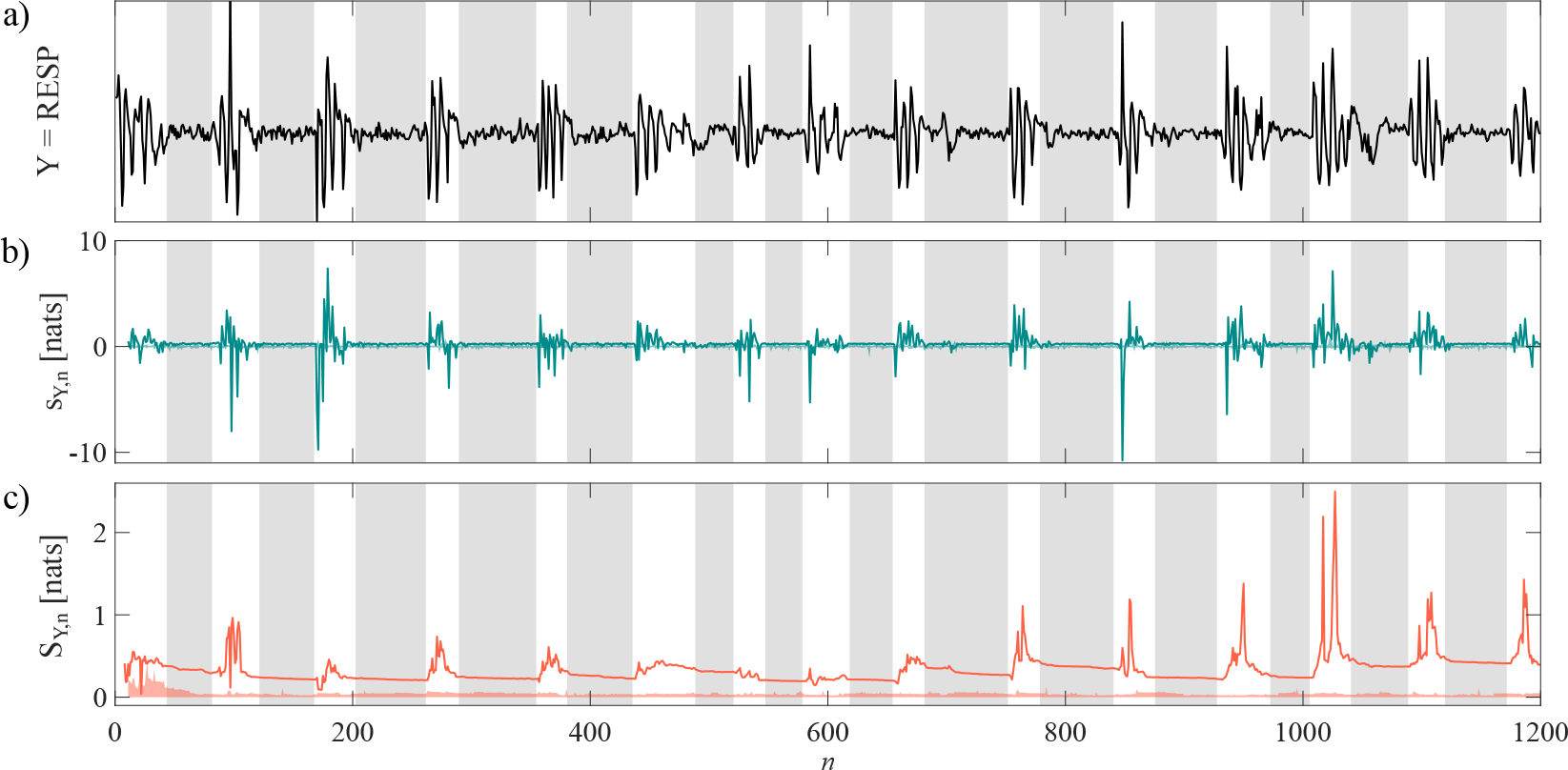
In panel *a*, respiratory time series (RESP) measured for a subject suffering of sleep apneas. Not-breathing intervals are evidenced using gray windows. Temporal profiles of the pointwise information storage measures obtained using local and time-varying approaches are depicted in panels *b* and *c*, respectively. Results obtained from one-hundred randomly time shuffled surrogate data are superimposed in transparency.

Results show how the L-IS profile takes statistically significant values only in the respiratory phases, with values above or below the surrogate distribution as this can take both positive and negative values. The TV-IS measure always exceeds the significance threshold, with values above the surrogate distribution regardless of respiratory dynamics which shows amplitude modulation with respiratory phases.

Figure 4 depicts the distributions of mean (panel *a*) and variance (panel *b*) of L-IS and TV-IS measured during apneas (A, gray boxplots) and breathing (B, white boxplots) intervals. A non-parametric Wilcoxon signed rank test for unpaired data (*α* = 0.05) was used to look at the differences of these indices during apneas (14 intervals) and respiration (15 intervals). Results highlight how both estimation approaches reveal strongly statistically significant difference in terms of variance computed in each interval (*p*-values in the range of 10^−6^), and the average values show a significant difference only for the time-varying approach (*p* = 0.0037).

**FIG. 4.**
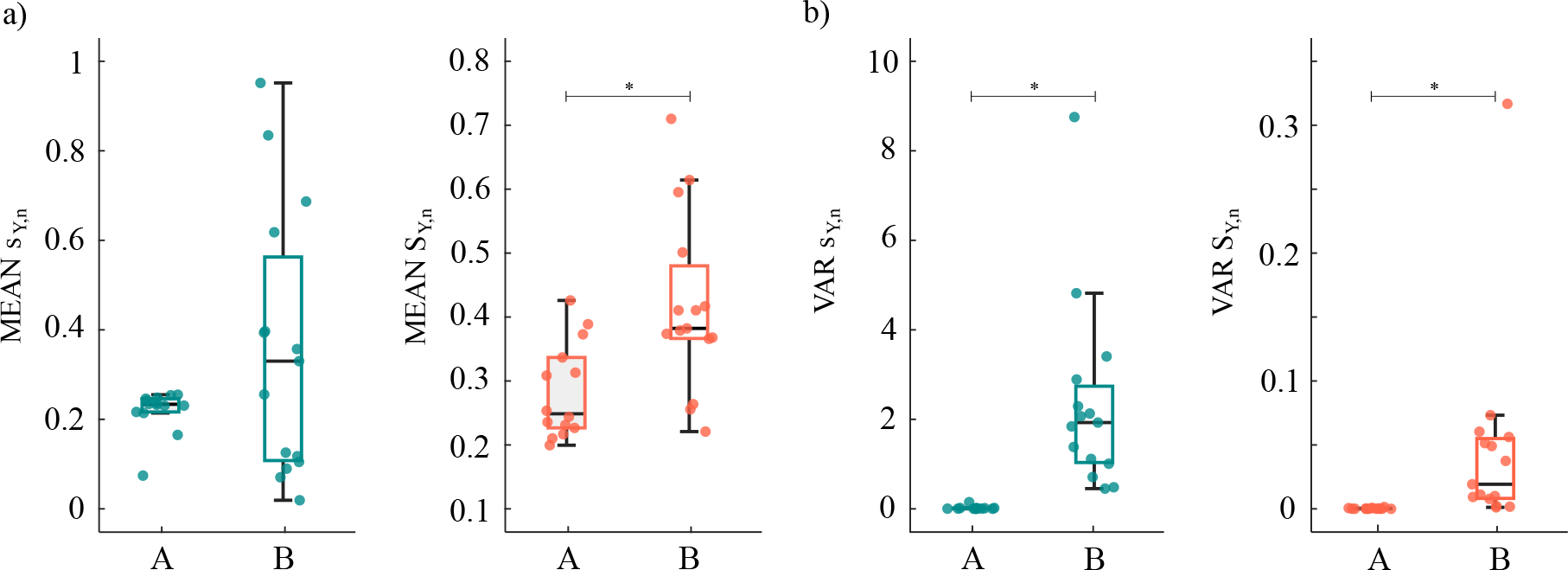
Boxplot distributions and individual values of L-IS and TV-IS representative of the mean (MEAN, panel *a*) and variance (VAR, panel *b*) in apneic (A, gray) and breathing (B, white) epochs. Statistically significant differences between non-apnea and apnea windows are marked with * (*p <* 0.05, Wilcoxon signed rank test).

The time-varying measure consistently exhibits higher mean values during breathing compared to the preceding apnea. In contrast, the local profile demonstrates positive and negative mean values with high variability during breathing periods, with no significant difference observed (*p* = 0.2849). Mirroring the results of the simulations, we interpret the increased mean of TV-IS and variance of L-IS observed in the breathing periods as an indication of stronger predictable dynamics of the respiratory signal localized in time during the non-apneic periods as a result of the reduced respiratory oscillations [16].

#### B. Epicranial multichannel EEG

Epicranial multichannel EEG traces from ten rats were used to investigate regularity of neural patterns during unilateral whisker stimulations [17]. In particular, the dataset comprises 15-channels epicranial EEG signals recorded at a sampling frequency of 2 kHz and bandpass filtered online between 1 and 500 Hz. The entire dataset can be found at https://doi.org/10.6084/m9.figshare.5909122.v1 and further details are reported in previous works [18, 23]. This dataset has been largely investigated in several works to evaluate the results from time-varying connectivity estimators, such as Granger causality [18], and partial directed coherence estimated through RLS [14] and with Kalman filter [23], as structural and functional pathways related to stimuli are well known and provided strong expectation about a specific configuration of functional connections between cortical areas.

We firstly analyzed the somatosensory evoked potentials (SEPs) obtained through a simple averaging procedure for the activity recorded over specific electrodes along the 50 stimuli contained in the dataset. The grandaverage analysis of the SEPs reported in Figure 5 highlights a maximum voltage peak over the primary sensory cortex contralateral (cS1, Electrode (E) 12, Figure 5*a*) which starts around 5 ms after the onset on the stimulation and then vanishing at around 25 ms [17, 18]. Moreover, cS1 has well defined structural connections to specific contralateral parietal (E14, Figure 5*a*) and frontal (E10, Figure 5*a*) sensory-motor regions which become active immediately after cS1 as confirmed from the analysis of SEPs reported in Figure 5*b*. Concerning the ipsilateral S1 (iS1, E4, Figure 5*a*), previous works suggested its involvement given the well known functional interactions between ipsi- and contralateral S1 [28, 29] and this is confirmed by the maximal activity measured over iS1.

**FIG. 5.**
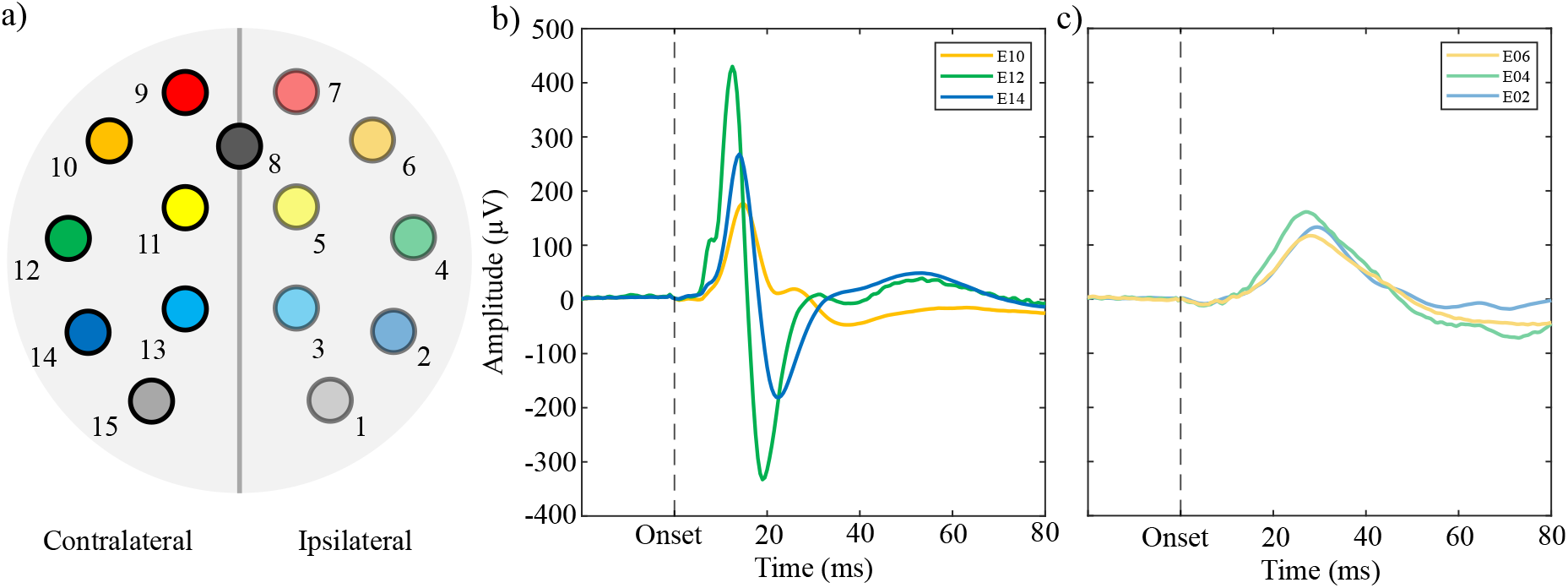
Schematic representation of electrode montage for epicranial recordings in panel *a*. Grand-average SEP for the contralateral (panel *b*) and ipsilateral (panel *c*) hemisphere to the whisker stimulation.

To study the modification over time of the information stored in the system, we considered data from the six electrodes relevant to the processing of the stimulus in the contralateral and ipsilateral hemisphere (i.e., E2,4,6,10,12 and 14). The entire recording comprising 50 trials of repeated stimulations was considered as a single realization of a stochastic process with a final time series length of *N* = 18000 time points. The time series was then reduced to zero mean and unit variance and the optimal model order was estimated for each electrode through the MDL criterion. The resulting model order was used for both L-IS and TV-IS estimation. Similarly to the previous application, the forgetting factor 1-*c* was set to 0.98 and the initial conditions of RLS (*A*_*p*_,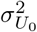) were obtained via the ordinary least squares prediction method. For each approach, the obtained estimates were then divided into 50 windows (one for each individual stimulus). As in the previous application, the mean and variance of the L-IS and TV-IS measured within the interval from onset to 40 ms after onset were calculated for each window. The values obtained were then averaged to obtain distributions relevant to the 10 animals.

The analysis of the results reported in Figure 6 reveals that both approaches can highlight a decrease of the predictability of the EEG signals associated with the onset of the SEP which is then restored after 25 ms. The drop of predictability is more evident in the cS1 (E12), where the SEP originates, and it is less prominent in the iS1 as expected from the physiology [17, 18].

**FIG. 6.**
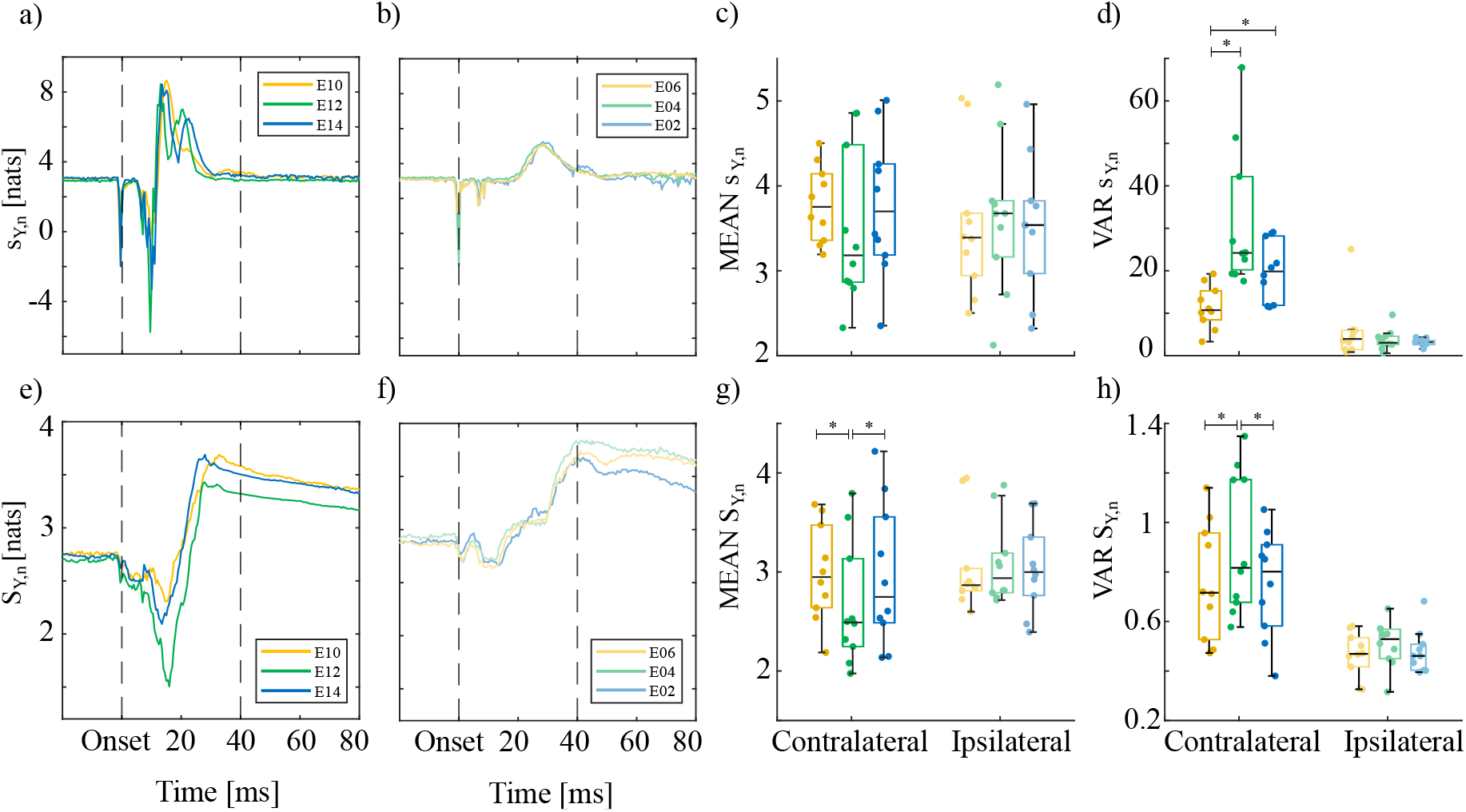
Grand-average distributions obtained for each of the six electrodes relevant to the processing of the whisker stimulation in the contralateral (panels *a*,*d*) and ipsilateral (panels *b, e*) hemisphere obtained by estimating local IS (top row) and time-varying IS (bottom row) of the EEG signals. Barplots are drawn evaluating mean (MEAN) and variance (VAR) of each approach (L-IS, panels *c-d* ;TV-IS, panels *g-h*) in the time interval [Onset, Onset+40 ms] marked by black dotted lines. Statistically significant differences between pairs of average values distributions are marked with * (*p <* 0.05, Wilcoxon signed rank test).

Boxplots depicting the distributions across animals of mean and variance of each approach highlights a statistically significant modulation of the variance of the L-IS in the contralateral primary sensory cortex (E12, panel *d*) which cannot be associated with increased mean values (panel *c*) as confirmed by the non-parametric Wilcoxon signed rank test (*α* = 0.05). Conversely, TV-IS shows a statistically significant decrease in the predictability occurring in cS1 (E12, panel *g*), with much lower variance values if compared to the L-IS. In the ipsilateral hemisphere, these differences are less prominent and not sta-tistically significant confirming the reduced relevance of the iS1 [18]. Reflecting the results of the simulation study and those obtained in the analysis of respiratory dynamics, we can explain the decreased mean of TV-IS and the increased variance of L-IS observed in cS1 during whisker stimulations as indicative of an increase in the complexity of the brain dynamics. Indeed, the whisker-evoked activity propagates from primary somatosensory cortex in the contralateral hemisphere which, in turn, is well known to represent the main hub in the motor network as evidenced in previous literature exploiting time-varying brain connectivity estimators to analyze the same dataset [18, 23].

## IV. CONCLUSIONS

In this study, we compared two approaches for detecting transient neurophysiological states, emphasizing their similarities and complementary aspects. To this end, we performed a simulation study specifically designed to reproduce transitions in the system dynamics, and then we applied the two approaches to benchmark data related to respiratory dynamics during transient sleep states and evoked responses in brain signals, where the underlying phenomena have been widely investigated.

The peculiarities of both approaches have been partially explored in previous works where the variability of local entropy measures was related to the surprise (the new information gained when an event occurs) induced by the emergence of a change in the system state [5, 10, 16], while the mean over specific time intervals of the TV-IS was related to the influence of sudden changes occurring locally in the internal dynamics of the brain system [11]. The extensive analysis performed in this paper confirms these properties and provides new insight about how pointwise and time-varying methods complement each other to reveal the time-resolved behavior of complex systems. Specifically, we conclude that: (i) the local predictability of a dynamic system can be influenced by both the evolution of its self-dependencies over time, which is directly related to the coupling between present and past states of the process, and by its unpredictable dynamics, quantified by the system noise; (ii) the mean of the TV-IS over specific time windows highlights the modulations of the predictable part of the system dynamics, while the variance of the L-IS measure reflects the modulation over time of both the predictable and unpredictable parts of the system dynamics; (iii) transient dynamics of neurophysiological systems can be described in terms of detectable changes in the mean (evidenced by the time-varying approach) or in the variability (evidenced by the local approach) of the pointwise information stored in the output signals of these systems.

Despite the potentialities of the two approaches, there are some limitations that should be taken into account. In our examples, we reasonably assumed the stationarity of the stochastic processes since multiple repetitions of the system’s oscillatory patterns were always present. However, when stationarity cannot be guaranteed, as in the case of single-trial analysis of SEPs, the local approach requires analysis across several realizations (trials). Since it is well acknowledged that physiological systems exhibit emergent behaviors which, so far, have been mostly studied under stationarity assumption, future developments will be focused on the exploitation of time-variant and local approaches to characterize transitions of higher-order interactions [30, 31]. This could have an impact on the study of altered and physiological states of different physiological systems, such as the cardiorespiratory and cardiovascular systems [32–35] along with the brain [36, 37], as well as of their interplay.

## ACKNOWLEDGMENTS

This work was supported by SiciliAn MicronanOTecH Research And Innovation CEnter “SAMOTHRACE” (MUR, PNRR-M4C2, ECS 00000022), spoke 3 - Universit`a degli Studi di Palermo “S2-COMMs - Micro and Nanotechnologies for Smart & Sustainable Communities. R.P. was partially supported by European Social Fund (ESF) Complementary Operational Programme (POC) 2014/2020 of the Sicily Region.

